# Dr.Paso: Drug response prediction and analysis system for oncology research

**DOI:** 10.1101/237727

**Authors:** Francisco Azuaje, Tony Kaoma, Céline Jeanty, Petr V. Nazarov, Arnaud Muller, Sang-Yoon Kim, Anna Golebiewska, Gunnar Dittmar, Simone P. Niclou

## Abstract

The prediction of anticancer drug response is crucial for achieving a more effective and precise treatment of patients. Models based on the analysis of large cell line collections have shown potential for investigating drug efficacy in a clinically-meaningful, cost-effective manner. Using data from thousands of cancer cell lines and drug response experiments, we propose a drug sensitivity prediction system based on a 47-gene expression profile, which was derived from an unbiased transcriptomic network analysis approach. The profile reflects the molecular activity of a diverse range of cancer-relevant processes and pathways. We validated our model using independent datasets and comparisons with published models. A high concordance between predicted and observed drug sensitivities was obtained, including additional validated predictions for four glioblastoma cell lines and four drugs. Our approach can accurately predict anti-cancer drug sensitivity and will enable further pre-clinical research. In the longer-term, it may benefit patient-oriented investigations and interventions.

## Introduction

The unbiased, large-scale prediction of anticancer drug activity using tumor-derived molecular data is crucial to deliver on the promise of a more personalized, precise treatment of cancer patients (Caponigro and Sellers, 2011; Ross and Wilson, 2014). The prediction of drug sensitivity based on the analysis of large collections of cell lines offers significant opportunities for investigating clinical efficacy in a biologically-meaningful, cost-effective manner (Geeleher et al., 2014; Goodspeed et al., 2016; Wilding and Bodmer, 2014). Computational models for predicting anticancer drug sensitivity can aid in the selection and prioritization of candidate compounds for pre-clinical research (Costello et al., 2014; Rees et al., 2016; Reinhold et al., 2012; Stetson et al., 2014).

Although cell line-based models may not fully recapitulate tumor biology, appropriately validated models may accelerate patient-oriented research, and have already shown potential to generate clinically-relevant predictions in different oncology domains. Such models may complement and in some cases offer an early substitute for *in vivo* models that tend to be expensive, time consuming and less scalable. In the short-term, this could enable the generation of novel biological hypotheses in the lab and, in the longer term, guide therapeutic decision-making in the clinic.

Over the past few years, the investigation of cell line-based computational models for anticancer drug sensitivity prediction has been accelerated by publicly-funded efforts of large research consortia (Barretina et al., 2012; Iorio et al., 2016b; Reinhold et al., 2012; Yang et al., 2013). In particular, the Cancer Cell Line Encyclopedia (CCLE) (Barretina et al., 2012) and the Genomics of Drug Sensitivity in Cancer (GDSC) (Garnett et al., 2012; Yang et al., 2013) projects represented significant steps forward for the oncology and pharmacogenomics research communities. These projects have generated and shared (untreated) molecular data from thousands of cancer cell lines and their accompanying treatment sensitivity measurements for hundreds of experimental and clinically-approved drugs. To date, computational models have mainly emphasized the application of different widely-investigated multivariable statistical and machine learning models, such as linear models and support vector machines, with various versions of feature selection methodologies (Dong et al., 2015; Haverty et al., 2016; Jang et al., 2014). Despite their potential for accurately predicting drug sensitivity across multiple types of cancer cell lines, less attention has been given to the investigation of biological importance of the proposed drug sensitivity markers, which have ranged from one to hundreds of gene-based features. Moreover, the majority of reported models have not been evaluated on independently generated datasets (Azuaje, 2017). Although different studies have tested the resulting prediction models on independent cell line datasets, e.g., models trained and tested on the GDSC and CCLE dataset respectively, there is a lack of studies that experimentally validate predicted anticancer sensitivity on independent biological samples, including cell lines that were not included in the training and initial testing datasets (Cortes-Ciriano et al., 2016; Gupta et al., 2016; Jang et al., 2014).

Here, we present Dr.Paso: Drug response prediction and analysis system for oncology research (Figure 1A). Dr.Paso predicts drug sensitivity responses based on the (baseline) expression patterns of 47 genes, which represented “hubs” in a pan-cancer transcriptomic network extracted from more than 1K cell lines and are substantially implicated in a diversity of cancer-relevant biological processes. A computational prediction model based on the multiple-linear regression of the 47-gene expression values measured in hundreds of cell lines provided both accurate and robust prediction performance. First, the model was trained and cross-validated on a (discovery) dataset consisting of more than 10K cell line-drug experiments for 24 (targeted and cytotoxic) drugs. Next, the resulting model was tested on a second, more recently-released, (validation) dataset comprising almost 10K cell line-drug experiments that included 16 drugs also found in the discovery dataset. Dr.Paso’s prediction performance is comparable to, and in some cases outperforms, previously published computational models. Motivated by these findings, Dr.Paso next predicted sensitivity scores for 4 glioblastoma (GBM) cell lines, including three (stem-like) cell lines that were not included in the discovery and validation datasets, against 24 drugs. We selected the top three drugs predicted as highly effective together with a drug predicted as lowly effective (negative control), and performed *in vitro* tests on the 4 cell lines. As in the case of the public datasets, the sensitivity scores estimated by Dr.Paso were highly concordant with the observed *in vitro* responses. To further facilitate research, we offer Dr.Paso through a Web-based interface that allows users to predict drug sensitivity scores for their own samples and expression data. The following sections will describe in detail these research phases, which are outlined in Figure 1B.

**Figure 1.**
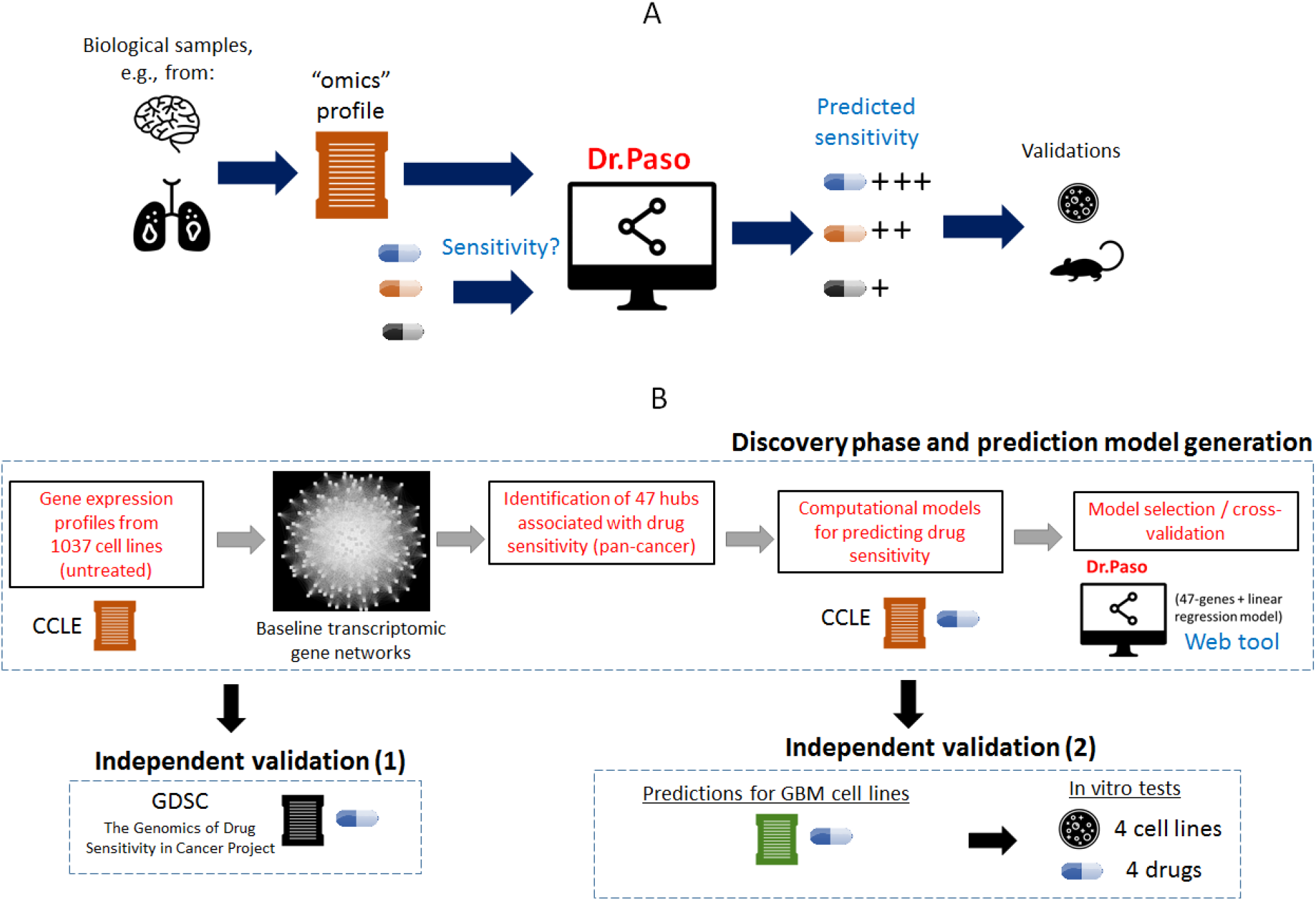
Dr.Paso: Overview of problem, application scenario and methodologies investigated. A. Outline of the general problem and application scenarios envisioned for the application of Dr.Paso. B. Workflow of the discovery phase, model generation and validation steps reported in this article.

## Results

### Hubs in a pan-cancer transcriptomic network display drug sensitivity predictive potential

Motivated by evidence indicating the drug sensitivity prediction power of gene expression profiles (Barretina et al., 2012; Iorio et al., 2016a), we investigated the predictive potential of such data in the context of a pan-cancer transcriptomic correlation network. Our hypothesis was that genes highly connected within such networks, i.e., hubs, may be reflective of molecular activity across biological processes and tissue sites. To test this hypothesis, we analyzed the CCLE gene expression dataset, which was derived from 1037 (untreated) cell lines representing different cancer types in 18 tissue sites. To reduce network complexity while aiming at preserving potentially relevant information across all samples, we selected genes with highly variable expression pattern across cell lines (i.e., 177 genes with standard deviation of expression values across cell lines located above the 99^th^ percentile). Using the pan-cancer expression profiles from these genes, we calculated all the between-gene (Pearson) correlation values and merged them into a fully-connected weighted network (Figure 2A), which included 177 nodes and more than 15K edges (correlations).

**Figure 2.**
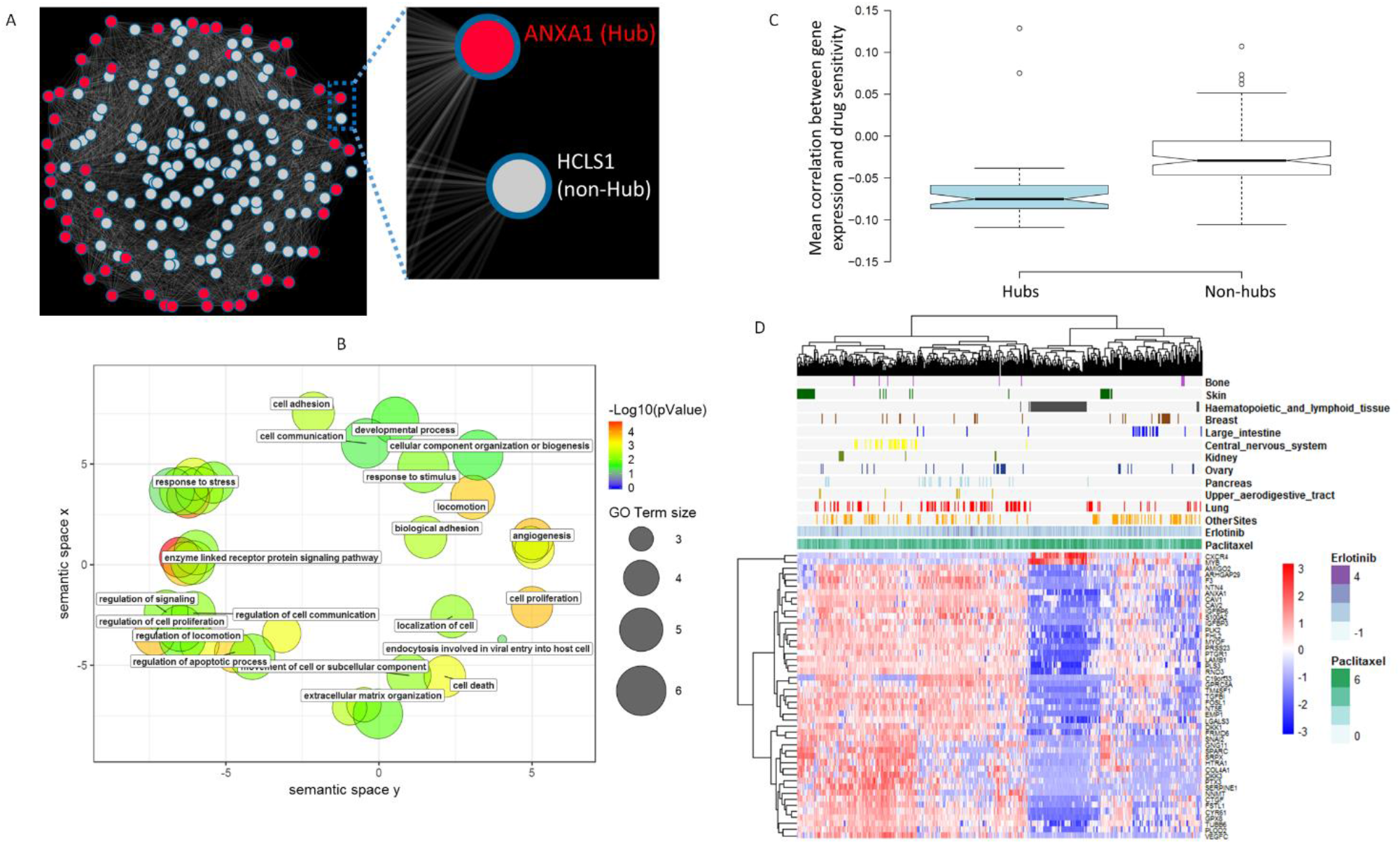
Hubs in a pan-cancer transcriptomic network display drug sensitivity predictive potential. A. Snapshot of a (fully connected) weighted gene correlation network. Nodes and edges representing genes and their correlations respectively. Network hubs and non-hubs are colored in red and white respectively. A zoom-in view of examples of hub and non-hub nodes. The color intensity of edges reflect the expression correlations between such nodes and others in the network. B. Graphical summary of (non-redundant) Gene Ontology terms statistically over-represented in the list of 47 hub genes. Terms are projected onto a semantic similarity space with REViGO (Supek et al., 2011), in which similar terms are positioned closer to each other. Each term is represented by a bubble with color and size indicating the term’s level of statistical enrichment in our list and frequency in the GO database respectively. C. Comparison of hubs vs. non-hubs on the basis of their individual associations with drug sensitivity. The boxplot depicts the mean correlation between the gene expression and the AA values across CCLE cell lines. Box notches indicate 95% confidence interval for each median value. D. Cell line-drug experiments are visualized in terms of the 47-gene expression data. The panel above the gene expression heatmap illustrates the AA values observed for selected sets of cancer cell lines (grouped by tissue site) and two compound examples (Erlotinib and Paclitaxel) for illustration purposes.

We identified network hubs by extracting those genes with statistically detectable connectivity scores (i.e., weighted degree values) using WiPer (Azuaje, 2014). This resulted in 47 hubs (WiPer adjusted-P < 0.05, online resource and Figure S1), one of which (*ANAX1*) is illustrated in Figure 2A together with an example of a non-hub node (*HCLS1*). The 47 hub genes are significantly implicated in a wide diversity of biological processes and pathways of relevance to cancer progression and therapeutic response. They include cell proliferation, death, migration, adhesion, angiogenesis, kinase signaling and the extracellular matrix (Figures 2B and S1).

Next, we analyzed drug sensitivity data (activity areas, AA) associated with these cell lines (11670 cell line-drug experiments) available in the CCLE. The AA, which is inversely correlated with the IC50 (the drug concentration at which an inhibition of 50% of cell viability is achieved), was defined by the CCLE to approximate the efficacy and potency of a drug simultaneously (Barretina et al., 2012). We stress that such data were not considered during the network generation and analysis steps. For each gene in the network, we calculated the correlation between gene expression and AA across all available (cell line-drug) data, and observed that: a. hubs tend to be anti-correlated with drug sensitivity, and b. such an anticorrelation is significantly stronger than in the case of non-hub genes. Moreover, such an association is considerably different to that displayed by non-hubs (Figure 2C). The 47 hub genes did not include previously reported markers of drug sensitivity: *ALK, BRAF, ERBB2, EGFR, HGF, NQ01, MDM2, MET and VEGFRs* (Barretina et al., 2012; Safikhani, 2017). To further demonstrate the potential relevance of these genes, we clustered the samples (available cell line-drug experiment data) based on their 47-gene (baseline) expression profiles and verified that these genes could, in principle, segregate samples according to cancer types (tissue sites) and highlight differential drug responses across samples (Figure 2D). Using an alternative visualization and (unsupervised) clustering technique (Figure S2), we verified the potential of these 47 genes’ expression data to segregate samples in terms of their drug sensitivity. Overall, these results suggest that our 47 hubs represented a potentially novel, biologically-meaningful gene set with drug sensitivity prediction potential.

### Predicting drug sensitivity based on the network-derived 47-gene expression profiles

We used the expression values from the 47 network hubs and drug sensitivity data (n = 10981, cell line-drug experiments, i.e., samples, with full expression and AA data available) to generate a drug sensitivity prediction model based on multiple linear regression (Methods). For a given sample (47-gene expression profile) and drug (identity of one of the 24 CCLE drugs), the model estimates a sensitivity score that approximates the AA values observed in the CCLE. For model training and testing, we used separate datasets respectively through a 10-fold cross-validation sampling procedure. Prediction capability was evaluated with multiple performance indicators that compare the predicted and observed sensitivity values: Pearson, Spearman and Kendall correlations, root-mean-squared errors (RMSE) and a concordance index (Figure 3).

**Figure 3.**
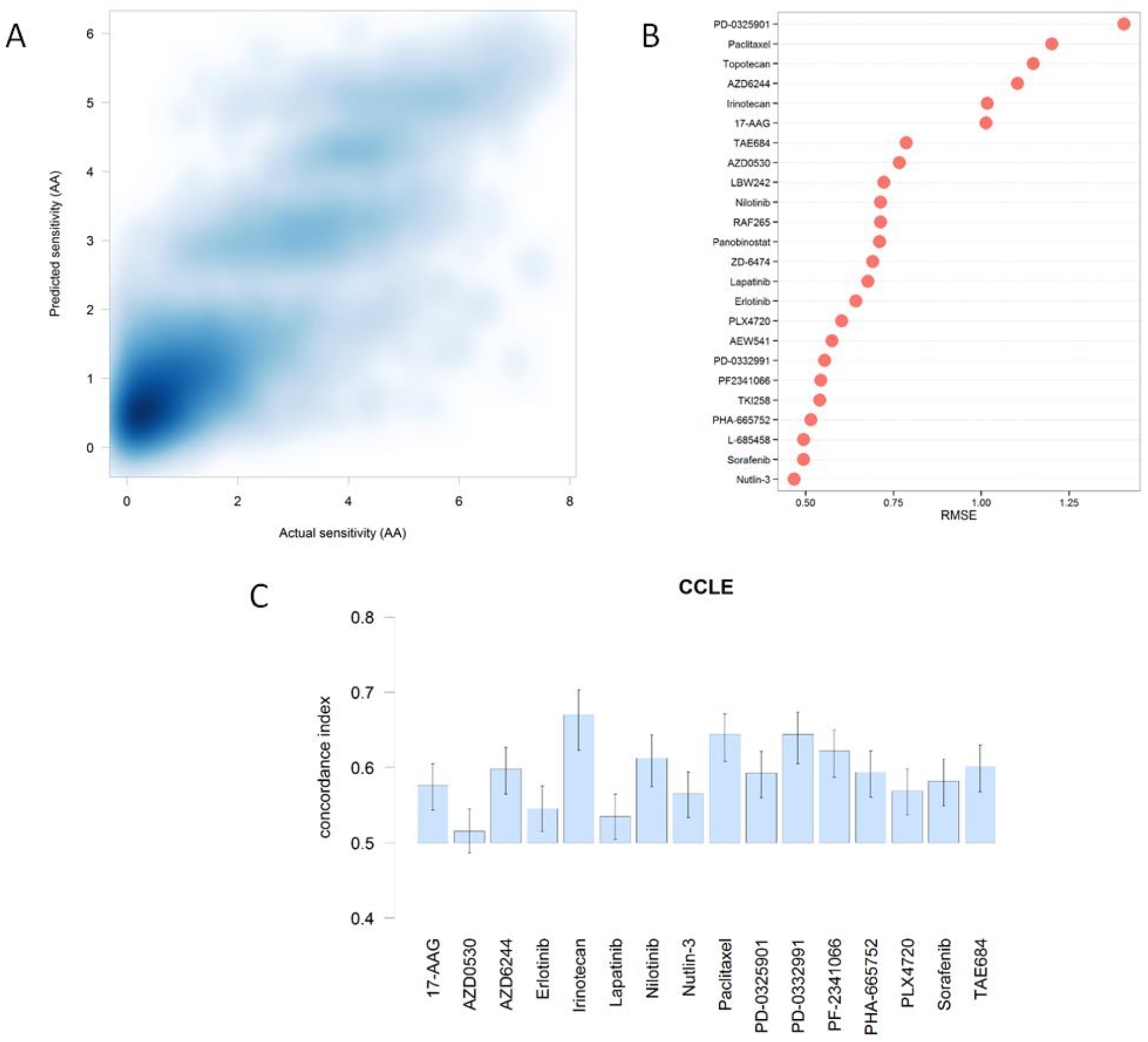
Alternative views of our model’s predictive capacity on the CCLE dataset using alternative performance indicators. A. Density plot of predicted vs. actual sensitivity values (n=10981). Pearson, Spearman and Kendall, correlations coefficients: 0.86, 0.73 and 0.54 respectively. B. Plot of root-mean-square errors (RMSE) observed for each drug. C. Concordance indices between the predicted and the observed AA values for a selected set of drugs. An index value = 0.5 is the expected value from random prediction. Error bars: 95% confidence interval (CI) of the estimated concordance index.

Figures 3A and S3 show that the predicted and actual AA values are positively correlated (Pearson, Spearman and Kendall, correlations coefficients: 0.86, 0.73 and 0.54 respectively), which corroborates the predictive potential of our model. Such performance measures are comparable to, and in many cases outperform, those obtained from other published models trained and cross-validated on the same dataset. For example, a comprehensive analysis of different machine learning techniques (Jang et al., 2014), e.g., multiple-linear regression techniques, support vector machines and random forests, displayed (median) Pearson correlation coefficients falling into the range from 0.4 to 0.6, including top-performing models generated with gene expression data or their combination with other data types (Jang et al., 2014).

Figure 3B offers an alternative assessment of our model’s prediction capability based on the RMSE obtained for each CCLE drug. This plot offers two key insights: 1. There are drugs for which our model can make relatively very accurate sensitivity predictions (e.g., Nutlin-3, an inhibitor of p53-Mdm2 complexes, and Sorafenib, a muti-kinase inhibitor) in comparison to other drugs (e.g., PD-0325901, a MEK inhibitor, and Paclitaxel, a mitotic inhibitor). 2. Our model’s (drug-specific) prediction performance is competitive in relation to other published approaches trained and tested on the same dataset. For example, our model made predictions with a median RMSE = 0.70 (range: [0.47, 1.40]), which compares well with top-performing machine learning models that have reported median and minimum RMSE values above 0.80 and 0.65 respectively (Neto et al., 2014). For drugs such as Sorafenib, Nutlin-3 and PHA-665752, Dr.Paso tends to outperform models based on elastic-net and other variations of multiple-linear regression (Neto et al., 2014). Conversely, such models tend to offer relatively more accurate predictions for drugs such as Irinotecan and PD-0325901. These results corroborate previous findings about the lack of generalized solutions for highly accurately predicting sensitivity across all types of drugs (Fersini et al., 2014; Haverty et al., 2016; Jang et al., 2014).

To provide further insights into our model’s prediction capacity, Figure 3C displays the concordance index for a selected set of drugs. For a random pair of samples, the concordance index estimates the probability of correctly predicting the relative sensitivities of such samples (e.g., sample X is more sensitive than sample Y) in relation to the actual observed relative sensitivities. Perfect and random prediction performances are indicated by concordance indices equal to 1 and 0.5 respectively. Our model reported concordance indices with median values above 0.5, which compares favorably with the results obtained by (Papillon-Cavanagh et al., 2013) with different alternative models, including multiple linear regression with elastic net, and applied to the same dataset. For instance, Papillon-Cavanagh et al. obtained concordance indices lower than 0.7, including predictions with concordance indices falling below 0.5 for different drugs (e.g., Nutlin-3 and TAE684). These results suggest that our model can accurately predict drug sensitivity and provide, in relation to previously published models, promising predictive capability that we further investigated as follows.

### Model evaluation on an independent dataset

We tested our 47-gene sensitivity prediction model on the 2016 release of the GDSC dataset (Iorio et al., 2016a). To allow our CCLE-derived model to make predictions on this dataset, we focused on the 16 drugs that are found in both datasets. First, as in the case of the CCLE data, we show that the (baseline) expression profiles of the 47 genes can, in principle, cluster the GDSC samples according to cancer types (tissue sites) and highlight differential drug responses across samples (Figure 4A) in an unsupervised manner. Note that in the GDSC dataset drug sensitivity is represented as the logarithm of IC50 (LNIC50) values (AA values were not provided in this dataset). Using an alternative visualization and (unsupervised) clustering technique (Figure S2), we verified the potential of these 47 genes’ expression data to segregate GDSC cell line-drug samples in terms of drug sensitivity.

**Figure 4.**
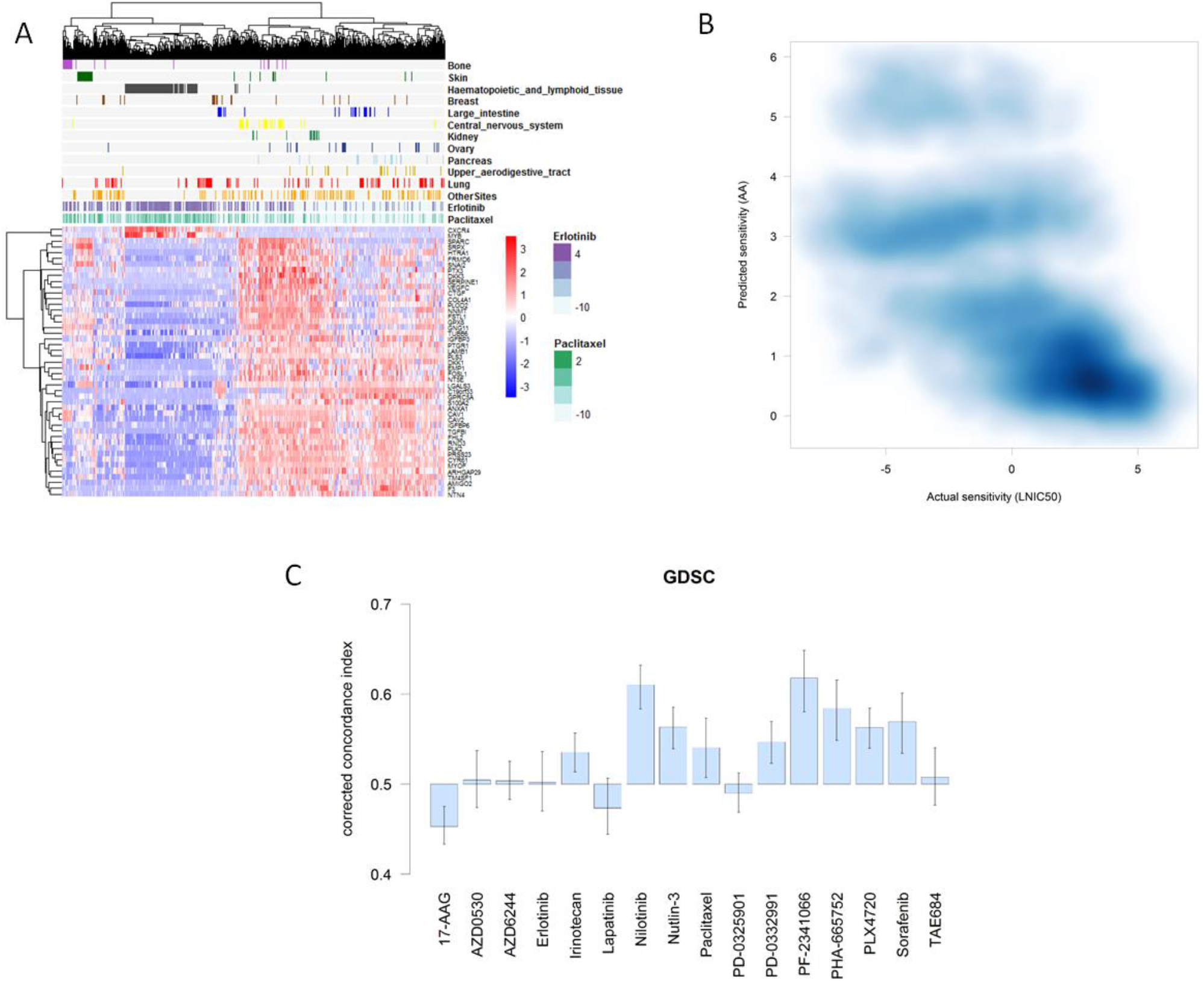
Alternative views of our model’s prediction capacity on the GDSC dataset. A. Cell line-drug experiments are visualized in terms of the 47-gene expression data. The panel above the gene expression heatmap illustrates the LNIC50 (μM) values observed for selected sets of cancer cell lines (grouped by tissue site) and two compounds (Erlotinib and Paclitaxel). B. Application of CCLE-derived model to the GDSC data. Density plot of predicted (AA) vs. actual sensitivity (LNIC50) values for drugs that are common between the CCLE and GDSC (n = 9984). Pearson, Spearman and Kendall, correlations coefficients: -0.72, -0.71 and -0.50 respectively. C. Concordance indices between the predicted and the observed sensitivity values. An index value = 0.5 is the expected value from random prediction. Indices are corrected to account for the notion that higher concordance is reached when high AA (prediction) corresponds to a low LNIC50 (observed) values, and vice versa. Error bars: 95% confidence interval (CI) of the estimated concordance index.

Next, we applied our (CCLE-derived) prediction model to the GDSC data and made sensitivity predictions (AA values) for all the samples (cell line-drug experiments) available (Methods). The resulting predictions were then compared with the actual sensitivity values in the GDSC dataset (Figures 4B and S3). The predicted (AA) and actual sensitivity (LNIC50) values for these samples (n = 9984) are anti-correlated (Pearson, Spearman and Kendall, correlations coefficients: -0.72, -0.71 and -0.50 respectively). This indicates that our model is, in general, estimating sensitivity values that are in agreement with those observed in the test dataset, i.e., higher predictive agreement is reached when high AA (prediction) relates to a low LNIC50 (actual) values, and vice versa.

Figure 4C summarizes the assessment of our model’s predictive performance on the GDSC dataset based on drug-specific concordance indices, as done for the CCLE dataset (Figure 3). Concordance indices > 0.5 were obtained for twelve out of the 16 drugs, and (among those 12 drugs) concordance estimates for 9 drugs can be reliably interpreted as larger than 0.5 (95% confidence intervals of the estimated indices). The predictive performances for several drugs (e.g., Nilotinib, Nutlin-3 and Sorafenib) are very similar to those estimated in the CCLE dataset. As in the CCLE dataset, the sensitivity observed in samples treated with AZD0530 and Lapatinib proved to be more difficult to accurately to predict. Although concordance indices > 0.5 were obtained for Irinotecan and Paclitaxel predictions, this represented a reduction of prediction performance in comparison to the predictions made for CCLE samples. The prediction performance of 17-AAG, PD-0325901 and TAE684 were also diminished. A previous study, which also used the GDSC dataset, consistently reported concordance indices < 0.5 for Sorafenib (Papillon-Cavanagh et al., 2013). Moreover, in comparison to that study’s models, our model reported comparable or higher concordance indices for other drugs, such as Nilotinib and PF-2341066 (Crizotinib). Conversely, such a previous study reported better prediction performances for 17-AAG, Lapatinib and PD-0325901. Such comparisons should, nevertheless, be interpreted with caution as Papillon-Cavanagh et al.’s concordance indices were obtained with an older version of the GDSC dataset, which was used for both model training and testing. Overall, our findings further corroborate the predictive potential of our model, and highlight strengths and challenges in a drug-specific context.

### Independent *in vitro* validation on several cell lines and compounds

To further validate our model’s predictive capability on independently-generated data, we generated predictions and performed *in vitro* tests for several GBM cell lines and compounds. First, we measured the (baseline) expression profiles of 4 GBM cell lines that have been well-characterized in our lab: U87, NCH644, NCH601 and NCH421k (Methods). While the CCLE and GDSC datasets included U87, the latter three are stem-like GBM cell lines that were not included in the previous model training and test phases.

Although genome-wide expression (microarray) data can appropriately cluster multiple samples (biological replicates) from such cell lines, we found that the expression profiles of our model’s 47 genes are sufficient to achieve the same biologically-meaningful segregation while offering a clearer, fine-grained view of their differences (Figure S4). We also verified the platform-independent replicability of these results with another 47-gene expression dataset derived from 3 of these cell lines measured with qPCR (Figure S4). These results corroborate the biologically-relevant discriminatory capacity and reproducibility of our model’s 47-gene expression patterns.

Next, our model predicted the sensitivity of our 4 GBM cell lines (18 samples in total, Methods) against the 24 drugs included in our model. The 47-gene (microarray) expression profiles of these cells were input to the prediction model (6 U87, 3 NCH644, 3 NCH601 and 6 NCH421k gene expression profiles). Figure 5A summarizes the 432 predicted sensitivity (AA) values according to drug (18 predictions per drug). To investigate such predictions *in vitro,* we focused on the top-3 drugs associated with the highest predicted sensitivities (Paclitaxel, Panobinostat and 17-AAG), as well as on Erlotinib, which was predicted as a relatively ineffective compound. These drugs correspond to four different drug classes: cytotoxic, histone deacetylase inhibitor, antibiotic derivative and an EGFR inhibitor respectively. In the case of Erlotinib, the predictions are consistent with the fact that the tested cells do not (NCH644, NCH421k) or very lowly (U87, NCH601) express EGFR. The Figures 5B and S5 show a more focused view of the predicted sensitivity values for our samples against these 4 drugs.

**Figure 5.**
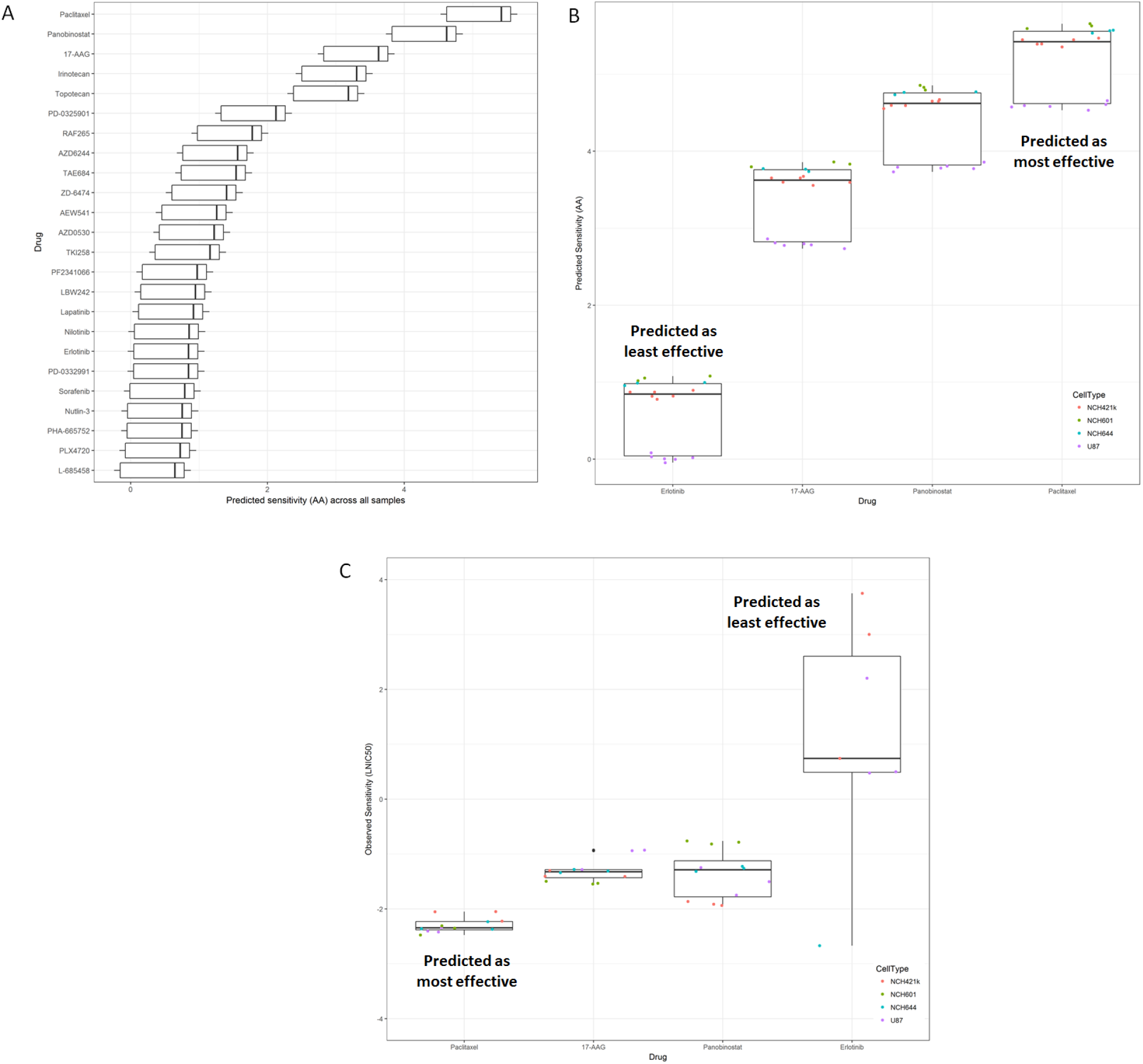
Drug sensitivity predictions and *in vitro* validation for different glioblastoma cell lines and compounds. A. Sensitivity predictions (horizontal axis) for 24 drugs (vertical axis). Box plot summarizes the (432) predicted sensitivity (AA, as defined in the prediction model) values for 4 glioblastoma cell lines: U87, NCH644, NCH601 and NCH421k. The 47-gene expression profiles of multiple biological replicates (18 samples in total) were input to the prediction model (6 U87, 3 NCH644, 3 NCH601 and 6 NCH421k samples). B. Alternative boxplot summary of the prediction results for 4 drugs (Erlotinib, 17-AAG, Panobinostat and Paclitaxel) and the different cell lines. These drugs, which were selected for subsequent *in vitro* tests, were predicted to be relatively highly (17-AAG, Panobinostat and Paclitaxel) and lowly (Erlotinib) effective against the 4 cell lines. C. Summary of *in vitro* test results. The selected drugs were tested on each cell line in triplicates, relative viability (vs. vehicle-treated samples) was measured for 8 drug concentration values (μM) and IC50 values were estimated for each drug-sample experiment. The boxplot shows the resulting LNIC50 values obtained. Drug response data for NCH601 samples and Erlotinib are not available, and for NCH644 samples and Erlotinib not shown because of lack of effect. Boxes show the median, the 25^th^ and 75^th^ percentiles (lower and upper hinges), and (1.5 x) inter-quartile ranges.

We tested the selected drugs on each cell line, in triplicates, and measured their response based on their relative viability (i.e., normalized to vehicle-treated samples) for 8 drug concentration values (μM). For each treated cell line, we estimated the IC50 values and compared them on the basis of cell line and drug groups. Figure 5C summarizes the results with boxplots showing the LNIC50 values. Drug response data for NCH601 samples and Erlotinib were not available (not tested), and data for NCH644 samples and Erlotinib are not shown due to lack of effect. Figure S6 includes all the drug response curves and additional details.

As predicted by our model, all our cell lines exhibited the lowest sensitivity, i.e., the highest IC50 values, when treated with Erlotinib (median LNIC50 = 0.74 μM). U87 was the least sensitive cell line in relation to all 4 drugs (median LNIC50 = -1.27 μM across all sample-drug experiments), in full agreement with the predictions. Our model consistently predicted NCH601 as the most sensitive cell line against all drugs (Figures S6). Our *in vitro* tests showed that NCH421k tended to be more sensitive than NCH601 (median logIC50: -1.64 vs. -1.54 μM). Despite this particular discrepancy, we found global agreement between predicted and observed sensitivities on the basis of cell type (Spearman correlation between the median sensitivity values, predicted (AA) vs. observed (LNIC50) in the 4 cell line groups: -0.40).

In accordance with the predictions, Paclitaxel was the most effective drug across all treated samples (median LNIC50 = -2.35 μM). Lesser agreement between predicted and observed sensitivities were obtained in the case of the remaining two drugs. For all samples, our model predicted overall higher sensitivity for Panobinostat than for 17-AAG (Figure 5B). *In vitro,* relatively similar responses were obtained for Panobinostat (median LNIC50 = -1.29) and 17-AAG (median LNIC50 = -1.33 μM), though a larger variability of sensitivity was observed in the former case. Nevertheless, predictions and *in vitro* tests concordantly showed that NCH421k and U87 samples treated with Panobinostat were consistently more sensitive than all samples treated with 17-AAG (Figures 5C and S6). Taken together, these results provide further evidence of the predictive capacity of our model. The resulting system, Dr.Paso, will enable the community to conduct further investigations.

### Dr.Paso online

To share our model and enable further research, we developed a web-accessible tool that allows researchers to upload their own gene expression data, make sensitivity predictions and visualize results in a few steps (Figure 6). The Help section of the website offers a guided application example using CCLE data. Users provide their input data as a text file containing the (baseline) 47-gene expression for different samples, and then can select all or specific drugs for making predictions (Figure 6A). Dataset re-scaling (feature standardization with means and standard deviations equal to 0 and 1 respectively) can be applied to harmonize the input dataset with the feature representation used in our model. Prediction results are presented with graphical displays and tables in different panels. Moreover, users can control the amount and focus of information at the drug and sample levels (Figure 6B to 6D). Results can be saved in different graphical and tabular file formats. The tool is freely available at www.drpaso.lu.

**Figure 6.**
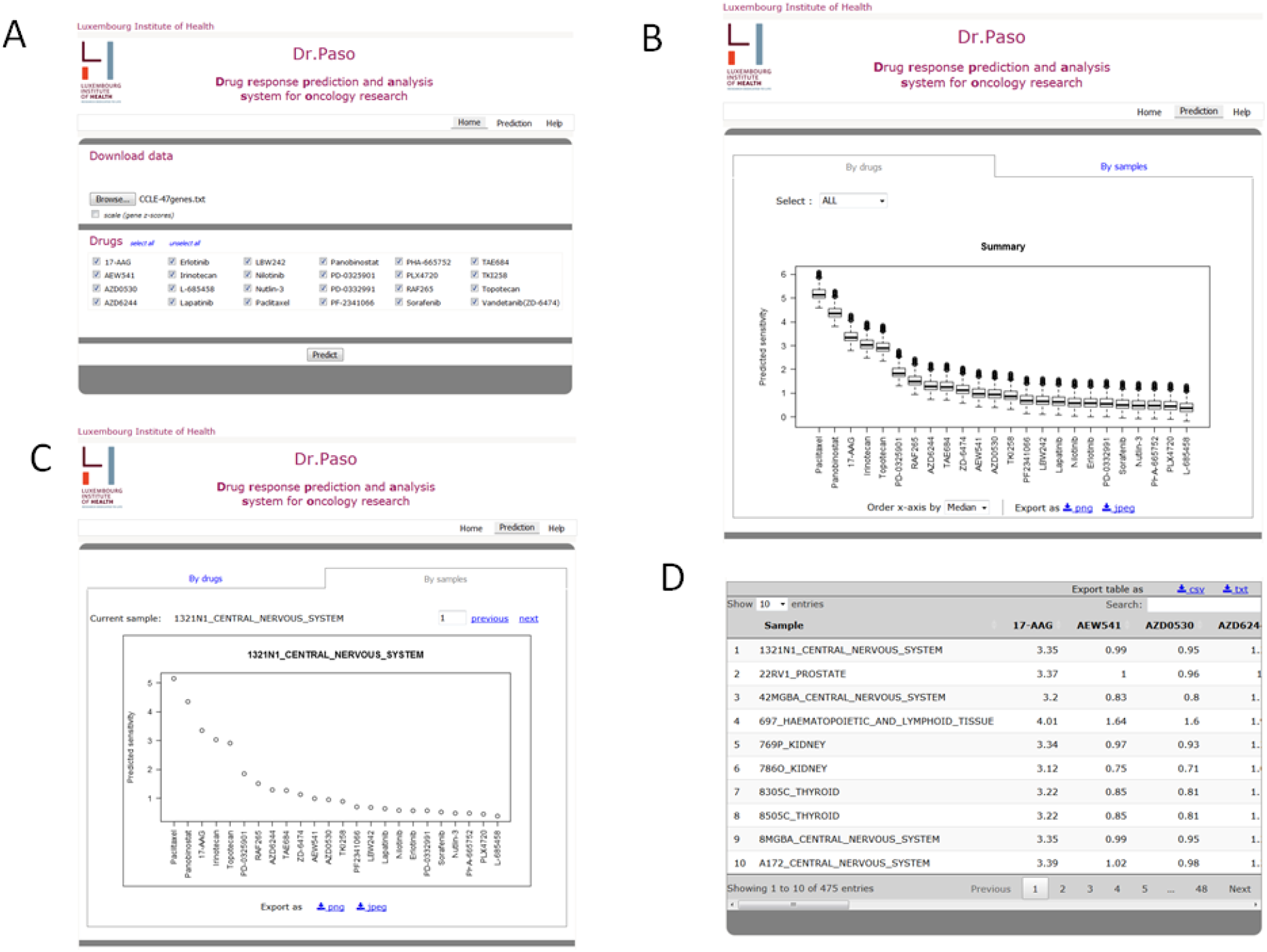
Dr.Paso online: a Web-based tool for predicting drug sensitivity and enabling further research. Screenshots of: A. Main page with user input and analysis options; B: Global view of predicted sensitivity values for a given input gene expression dataset and all drugs available in the CCLE; C: Alternative view of predictions focused on a specific input sample and all drugs; D. Tabular-based view of results. All views can be selected and downloaded according to user requirements.

## Discussion

The development of computational models for estimating drug sensitivity based on the analysis of large and diverse collections of cancer cell lines is important to support pre-clinical research, and provides a basis for future clinically-oriented applications. Access to such models and their user-friendly application will enable new research across oncology domains and additional computational investigations. Our Drug Response Prediction and Analysis System for Oncology research, Dr.Paso, addresses such needs through the integration of network-based and statistical modeling approaches. For a given drug, our system predicts an anti-cancer sensitivity score based on the gene expression profile of 47 genes, which were shown to represent hubs in a pan-cancer transcriptomic network and to be prominently implicated in a variety of cancer-relevant biological processes. Dr.Paso was generated and evaluated on independent datasets, including our own *in vitro* validations of several cell line-compound combinations, and showed promising results in terms of predictive accuracy and concordance. Future research can apply Dr.Paso to 47-gene expression signatures from patient samples to investigate its potential relevance in the clinical setting.

Our study and other previous research highlight the challenges faced and complementary predictive capacity exhibited by different modeling approaches (Costello et al., 2014; Papillon-Cavanagh et al., 2013). No single model can consistently make accurate predictions for all drugs and cell lines available in the CCLE and GDSC datasets, including models that include genomic data (Gupta et al., 2016; Jang et al., 2014; Menden et al., 2013). Different models can offer more, or less, accurate predictions for certain drugs, and there is no conclusive evidence about the dominance of a particular modeling technique (Azuaje, 2017). Such limitations may be partially explained by a lack of sufficient molecular information to account for the complexity of cell lines and their drug responses, choice of surrogate measures of drug sensitivity and inconsistencies of sensitivity data between the CCLE and GDSC (Haverty et al., 2016; Investigators, 2015; Safikhani, 2017). The latter may also partly explain the overall degradation of predictive performance when training models on the CCLE and testing them on the GDSC.

Dr.Paso generates sensitivity scores based on a multiple linear regression model. We, as others elsewhere, have shown that relatively less complex regression models can offer comparable, and in some cases better, prediction performance than those models based on larger sets of learning parameters. Dr.Paso’s predictive capacity is grounded in an unbiased network-guided selection of model inputs (47 genes) prior to the fitting of the regression model. Such a discovery process was shown to be both statistically- and biologically-meaningful. Apart from our multiple linear regression, we applied other regression techniques, e.g., support vector machines and neural networks, but decided to focus our investigation on a model with relatively lower complexity. Collectively, Dr.Paso is based on a biomarker discovery and prediction-making methodology that is both biologically-driven and statistically-powerful. New investigations, motivated by new datasets and clinically-oriented questions, are certainly envisaged and are expected to include new biomarker discovery and prediction modeling strategies.

Notwithstanding recent advances in the field, there is a need to make executable models accessible to the research community to enable new investigations, including new applications and comparative analyses among different techniques. Here we offer Dr.Paso as a publicly-accessible online tool. While further investigations are needed, our study offers further evidence of the potential of computational models for predicting anti-cancer sensitivity. In the short-term, our findings will enable new pre-clinical research applications and may provide a new perspective for bringing such models closer to the clinic.

## Methods

### Identification of 47 genes with drug sensitivity predictive potential

The published pre-processed CCLE (microarray) gene expression and drug sensitivity datasets were obtained from the CCLE website. In the gene expression dataset, we focused on genes with symbols, calculated their standard deviation (SD) across all samples (1037) and ranked them based on their SD. For further analyses, we selected the most variable genes: 177 genes with SD values above the 99^th^ percentile of the SD value distribution. We computed the gene-gene (Pearson) correlation coefficients between all the 177 genes and merged them into a single gene expression correlation network. We applied WiPer (Azuaje, 2014) to this fully-connected weighted network to detect highly connected nodes (hub genes). For each network node, WiPer computes the weighted degree and a corresponding P-value to assess the significance of the observed values, and adjusts it for multiple testing. Genes exhibiting (Bonferroni adjusted) P< 0.05 were considered hubs (47 genes). Drug sensitivity information was not used to select hubs. The resulting 47 genes were examined with different Gene Ontology (GO) and biological pathway analysis tools (below). For each hub gene, we estimated the correlation of its expression profile (across all samples) with the activity area (AA) values available from all sample-drug combinations. The AA was formulated by the CCLE to approximate the efficacy and potency of a drug simultaneously and is inversely correlated with the IC50 (Barretina et al., 2012). We compared hubs and non-hubs on the basis of such individual expression-sensitivity correlations. Visualizations and unsupervised clustering of hubs and cell lines described by hub expression values were implemented with different open-source tools (below).

### Training and testing of prediction model

We represented each CCLE sample (cell line-drug combination) with the expression values of the 47 hub genes and their corresponding AA values. We focused on samples with complete expression and AA data. The resulting set of 10981 samples was used for training and testing regression models. The dataset was standardized by re-scaling each gene so that each gene has mean and standard deviation of 0 and 1 respectively. For each model, we implemented 10-fold cross-validation (CV) for separating training from testing and for assessing prediction performance. We also used leave-one-out CV (LOOCV) and similar prediction performance results were obtained. Diverse regression techniques with different levels of complexity were investigated. We focused on a multiple linear regression model with Ridge regularization (Ridge parameter = 1E-08) because its performance (regression errors) was better than or comparable to those obtained with other techniques, such as support vector machines and k-nearest neighbors, and because of its interpretability in comparison to relatively more complex models. The accuracy of model predictions was assessed by measuring their (Pearson, Spearman and Kendall) correlations with the observed values in the CCLE, the root-mean-squared error (RMSE) and a concordance index. The latter approximates, for a random pair of samples, the probability of correctly predicting which sample is more (or less) sensitivity than the other (Harrell et al., 1996). A concordance index equal to 0.5 indicates that the model’s performance is comparable to that from a random predictor, while an index equal to 1 represents the perfect predictor.

### Independent evaluation on the GDSC dataset

Raw expression data were obtained from the ArrayExpress database (accession number E-MTAB-3610) and drug sensitivity (natural logarithm of the IC50 in μM, LNIC50) were downloaded from GDSC database (http://www.cancerrxgene.org, release-5.0). We normalized raw expression data with the RMA function of R/oligo package (Carvalho and Irizarry, 2010). Then we averaged the resulting log2 probe-set intensities to estimate the expression of each gene. Associations between probe-sets and gene symbols were obtained through the hgu219.db annotation package (Carlson, 2016). For each cell line-drug experiment available (sample), we retrieved the expression data for the 47 genes used as inputs to our prediction model and retrieved the corresponding drug sensitivity. We focused on the 16 drugs found in both this and the CCLE dataset. This resulted in a dataset consisting of 9984 samples, each one represented by 47 gene expression values and one LNIC50 value. We standardized expression data as in the case of the CCLE dataset, reformatted the file and input it to the CCLE-derived prediction model (further information below). For each sample in the dataset, the model predicted a drug sensitivity score (approximation of AA). We compared predicted vs. observed values using the indicators applied to the CCLE dataset analysis. We adapted the concordance index to account for the fact that AA and LNIC50 are expected to be anti-correlated, i.e., for a given sample, concordance is achieved when a high (predicted) AA value corresponds to a low (observed) LNIC50 value, and vice versa.

### GBM cell lines and expression data for *in vitro* validations

U87 cells were obtained from the ATCC (Rockville, USA) and were cultured as monolayers in DMEM containing 10% FBS, 2 mM L-Glutamine and 100 U/ml Pen-Strep (Lonza). GBM stem-like cultures (NCH421k, NCH601 and NCH644) were kindly provided by Christel Herold-Mende (University of Heidelberg, Germany) and were cultured as 3D non-adherent spheres as previously described (Abdul Rahim et al., 2017; Sanzey et al., 2015).

We measured the (baseline) gene expression of 4 GBM cell lines using microarrays (6 U87, 6 NCH421k, 3 NCH644 and 3 NCH601 biological replicates), as reported in (Sanzey et al., 2015). For our model’s 47 genes, we also replicated gene expression measurements using qPCR for U87, NCH421 k and NCH644 cell lines (each one in triplicate). RNA was extracted from 10^6^ cells using TRI Reagent^®^ (Sigma-Aldrich). RNA isolated in the aqueous phase with a Phase lock gel-Heavy (5 Prime) was precipitated with 100% isopropanol and purified using RNeasy^®^ Mini kit combined with an on-column DNase treatment (Qiagen). For the qPCR, RNA was reverse-transcribed into cDNA using Superscript III™ (Invitrogen) following manufacturer’s instructions. qPCR was performed in 96-well plates using SYBR^®^ Green Master Mix (Bio-Rad) and CFX-96 thermal cycler (Bio-Rad). Normalized gene expression levels were calculated using the CFX manager 3.1 software (Bio-Rad) via the delta-delta Cq method with “Hspcb, Rps13, 18sRNA” as reference genes and taking into account the calculated amplification efficiency for each primers pair. We provide a MIQE-compliance checklist table as a supplemental item.

### Drug sensitivity predictions and *in vitro* validation on GBM cell lines

The gene (microarray) expression dataset was standardized as above. Each sample, represented by a 47-gene (microarray) expression profile, was input to the prediction model and a drug sensitivity value was predicted for each one of them (18 samples in total), for each of the 24 drugs included in the model. Predicted values were compared between them to determine their relative differences in terms of cell lines and drugs. Next, these predictions were compared to the *in vitro* sensitivity values that were obtained as follows. We tested 4 drugs: Paclitaxel (Sigma-Aldrich), Panobinostat, 17-AAG and Erlotinib (Selleck Chemicals) independently on the selected 4 GBM cell lines with 8 drug concentrations. For each cell line and dose, we performed treatment experiments in triplicate (i.e., 3 treated biological replicates / dose). As a measurement of drug sensitivity, WST-1 (Sigma-Aldrich) cell viability assays were implemented. U87, NCH421k, NCH644 and NCH601 cell lines were seeded into 96-well plates at densities of 1,500, 5,000, 4,000, 6,000 cells per well, in appropriate culture medium (Sanzey et al., 2015). Cells were incubated, 24h hours after seeding, with the 8 different drug concentrations ranging from 10μM to 6.1 x 10-4 μM, with a final volume of DMSO not exceeding 0.1% and each condition was tested with 6 technical replicates. After 72h incubation, WST-1 reagent was added in medium to a final concentration of 10%. Adherent cell line (U87) was incubated at 37°C for 2 hours and 3D sphere stem-like cell lines (NCH421 k, NCH644 and NCH601) were incubated at 37°C for 6-8 hours. Absorbance was measured against a background control at 450nm on a FLUOstar OPTIMA Microplate Reader (BMG LABTECH). Using the normalized viability measurements, we generated drug dose-response curves and estimated IC50 values (μM) for each sample-drug combination. The dose-response curves were fitted with a four-parameter logistic regression model, whose parameters were calculated using GraphPad Prism 7 (GraphPad).

### Software and Dr.Paso’s Web-based tool

We used the R statistical environment for data analysis and visualization (www.r-project.org), packages: ggplot2, pheatmap, MASS and SNFtool (Wang et al., 2014). Concordance indexes (Harrell et al., 1996) were calculated based on rescaled Kendall rank correlation coefficients, which were also used to estimate confidence intervals (by Fisher’s transformation). For network analyses, we applied Cytoscape for visualization (Shannon et al., 2003), MINE for similarity exploration (Reshef et al., 2011) and WiPer for network hub identification (Azuaje, 2014). REViGO (Supek et al., 2011) and g:Profiler (Reimand et al., 2007) were applied for biological process and pathway enrichment analyses. The Weka workbench was used for building and testing regression models (Frank, 2016; Hall, 2009), and GraphPad Prism (www.graphpad.com) for analyzing drug response curves. We provide researchers with a Web-based application to enable them to predict anticancer drug sensitivity using their own (47-gene) transcriptomic data. The tool is based on the R/Shiny package (https://shiny.rstudio.com/). Although this package offers useful functionality for generating an interactive user interface, we customized available code using the R/Shinyjs package (http://deanattali.com/shinyis/). Users can input pre-processed expression datasets. Alternatively, our application can also implement z-score rescaling of the input data. Figures containing the prediction results can be downloaded and stored as either png or jpeg files. Results are also shown as tables with sample-specific predictions (in rows) with their corresponding drugs (in columns), and may be stored as either csv or tsv files.

## Author Contributions

Conceptualization, F.A.; Methodology, F.A., P.V.N, C.J., T.K., A.G, S.P.N; Software, F.A., T.K., P.V.N, C.J., A.M., S.K.; Validation, T.K, C.J.; Resources, A.G, G.D, S.P.N, Supervision, F.A., A.G., S.P.N, G.D.; Writing - Original Draft, F.A.; Writing - Review & Editing, all authors.

## Acknowledgments

This research was funded by (LIH-MESR) project Connect2Predict to F.A. For technical guidance to C.J, we thank H. Erasimus and S. Fritah (drug experiments) and V. Barthelemy, A. Bernard, J. Bohler and A. Dirkse (cell line manipulation) at the LIH NorLux NeuroOncology Laboratory. For helpful feedback on the manuscript, we thank L.C. Tranchevent at the LIH Proteome and Genome Research Unit.

## Supplemental Information

**Figure S1.**
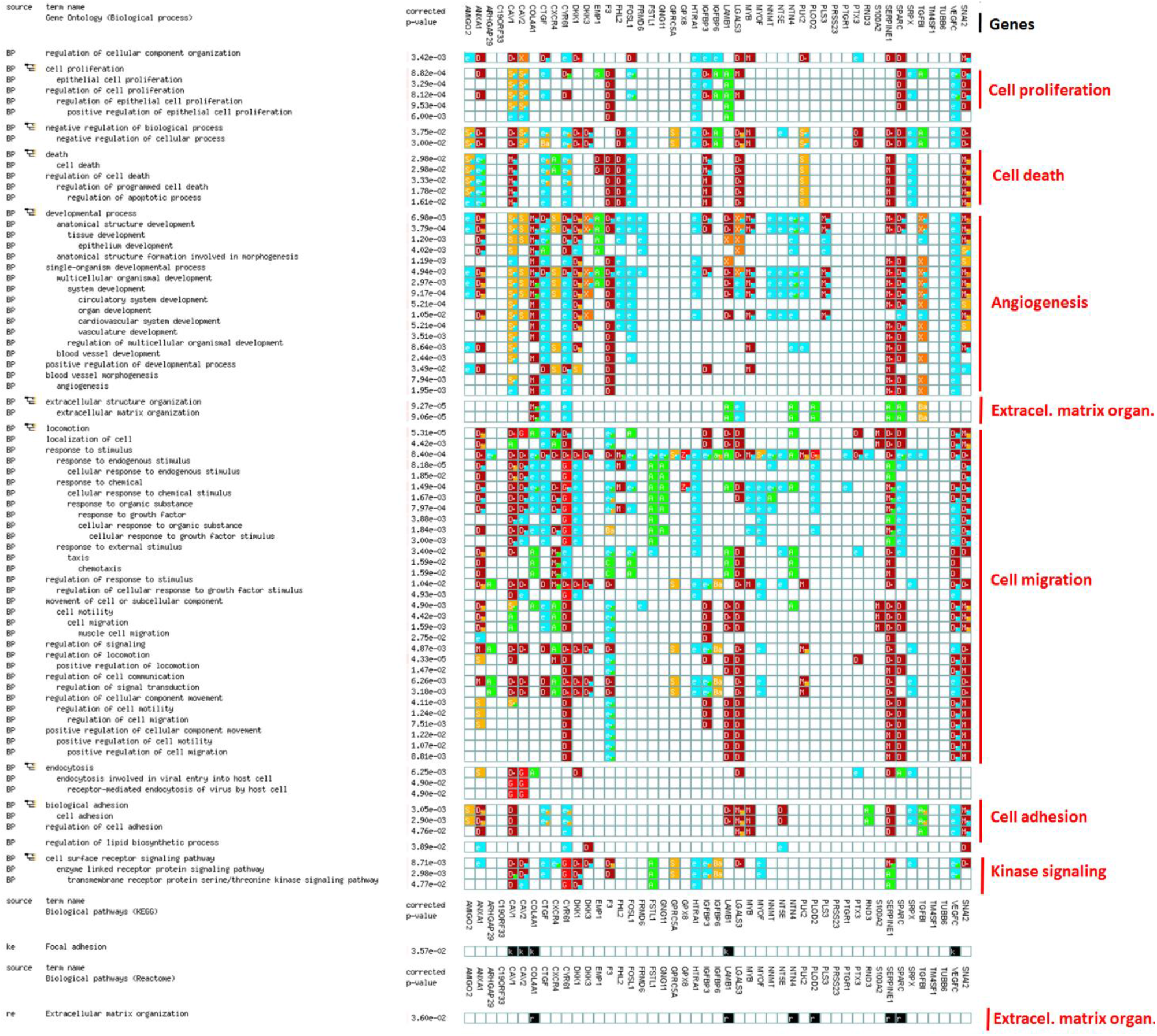
Statistical enrichment analysis of biological processes and pathways in the set of 47 network hubs. Related to Figure 2B. An alternative visualization of functional enrichments using G:Profiler (Reimand et al., 2007). As shown in Figure 2B, our set of 47 hub genes (columns) is significantly associated (at corrected P-value = 0.05) with a diversity of biological processes and pathways (rows). Each colored cell represents the association between an individual gene and a functional annotation. Colors are used to specify the evidence type of the observed association: 9: Inferred from experiment, Ell¡l: Direct assay / Mutant phenotype, Traceable author, : Electronic annotation, additional information at http://biit.cs.ut.ee/gprofiler.

**Figure S2.**
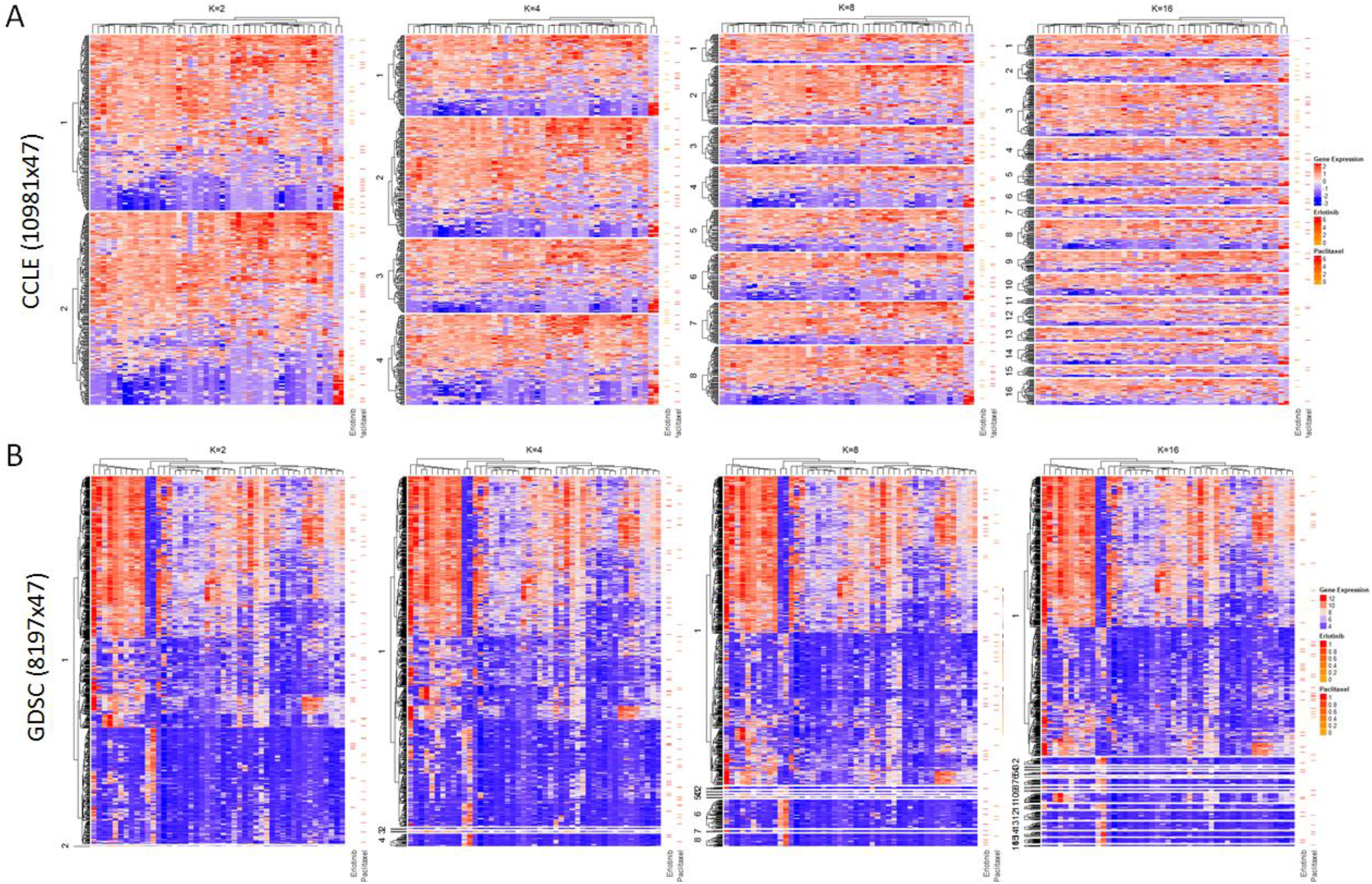
Alternative visualizations and unsupervised clustering of CCLE and GDSC cell lines on the basis of their 47-gene profiles. Related to Figures 2D and 4A. Spectral clustering analysis was applied using the SNFtool (Wang et al., 2014) to independently explore the potential of the 47 genes’ expression data to segregate (cell line-drug experiment) samples. A. CCLE and B. GDSC results. In A. and B., rows and columns in each heatmap represent samples and genes respectively, and color represents gene expression intensity. To facilitate visualization, clustering results for different numbers of clusters (K) are provided as independent plots. Note that the order of the rows in each clustering (plot) is not preserved. In each plot, additional columns (right side) representing the drug sensitivity of the samples against Erlotinib ad Paclitaxel are illustrated.

**Figure S3.**
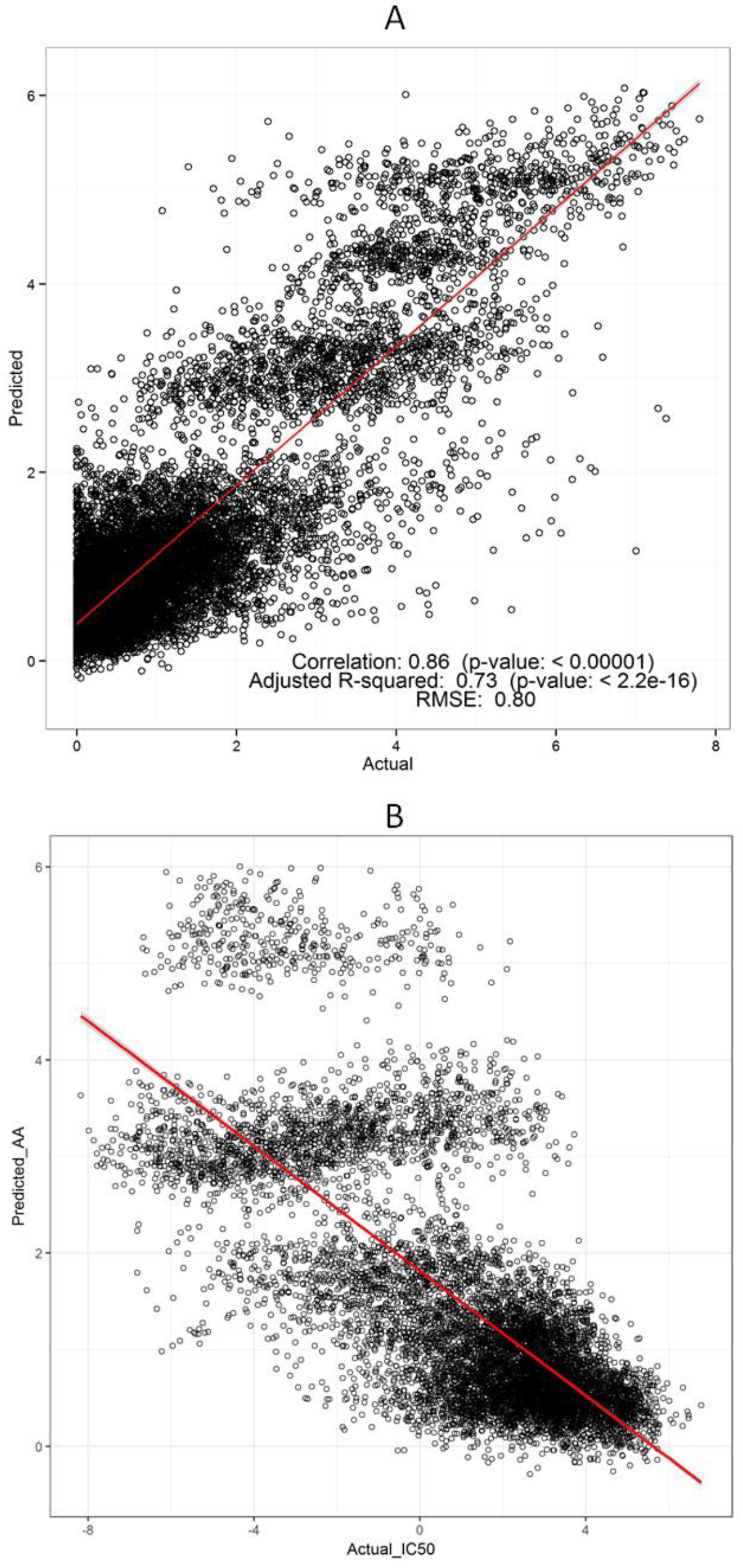
Predicted vs. actual sensitivity values in the CCLE and GDSC datasets Related to Figures 3A and 4B. Alternative visualization to those shown in Figure 3A. A. CCLE plot (n=10981). B. GDSC plot (n=9984).

**Figure S4.**
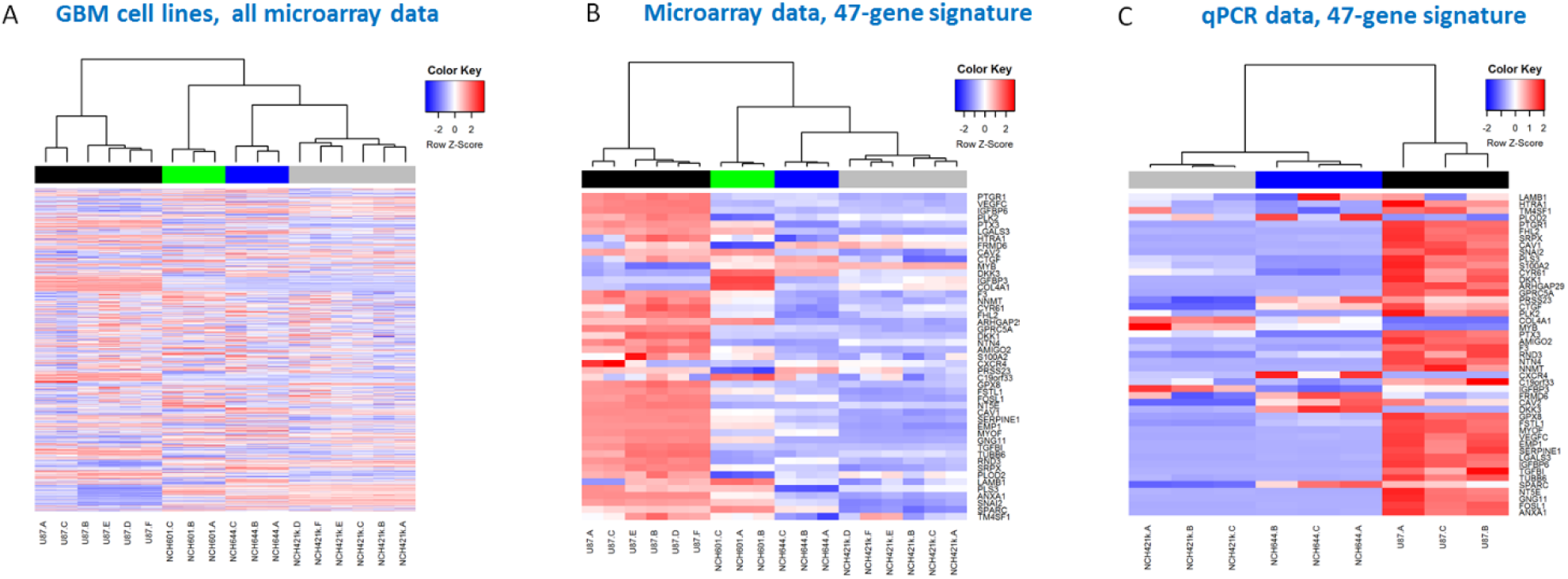
The 47-gene signature distinguishes cell types and is reproducible. Related to section: “Independent *in vitro* validation on several cell lines and compounds”. Gene expression of 47 genes in 3 GBM cell lines using microarrays and qPCR. Analysis peformed to verify the robustness and platform-independent replicability of the 47-gene expression data and its capacitiy to distinguish between cell lines.

**Figure S5.**
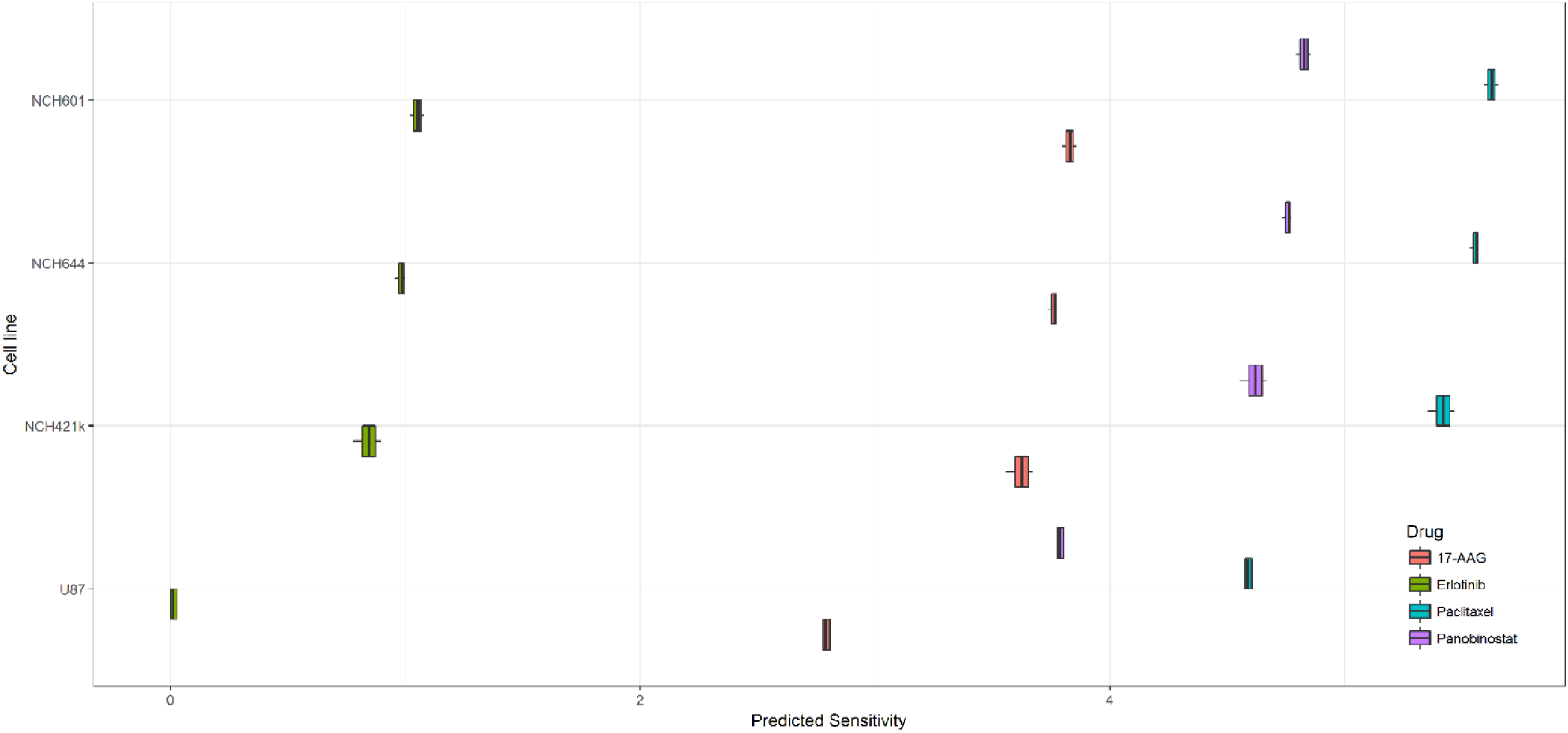
Boxplot summary of prediction results for 4 drugs (Erlotinib, 17-AAG, Panobinostat and Paclitaxel) and 4 GBM cell lines. Related to Figure 5B. Ech cell line type comprises multiple biological replicates (18 samples in total): 6 U87, 3 NCH644, 3 NCH601 and 6 NCH421 k samples.

**Figure S6.**
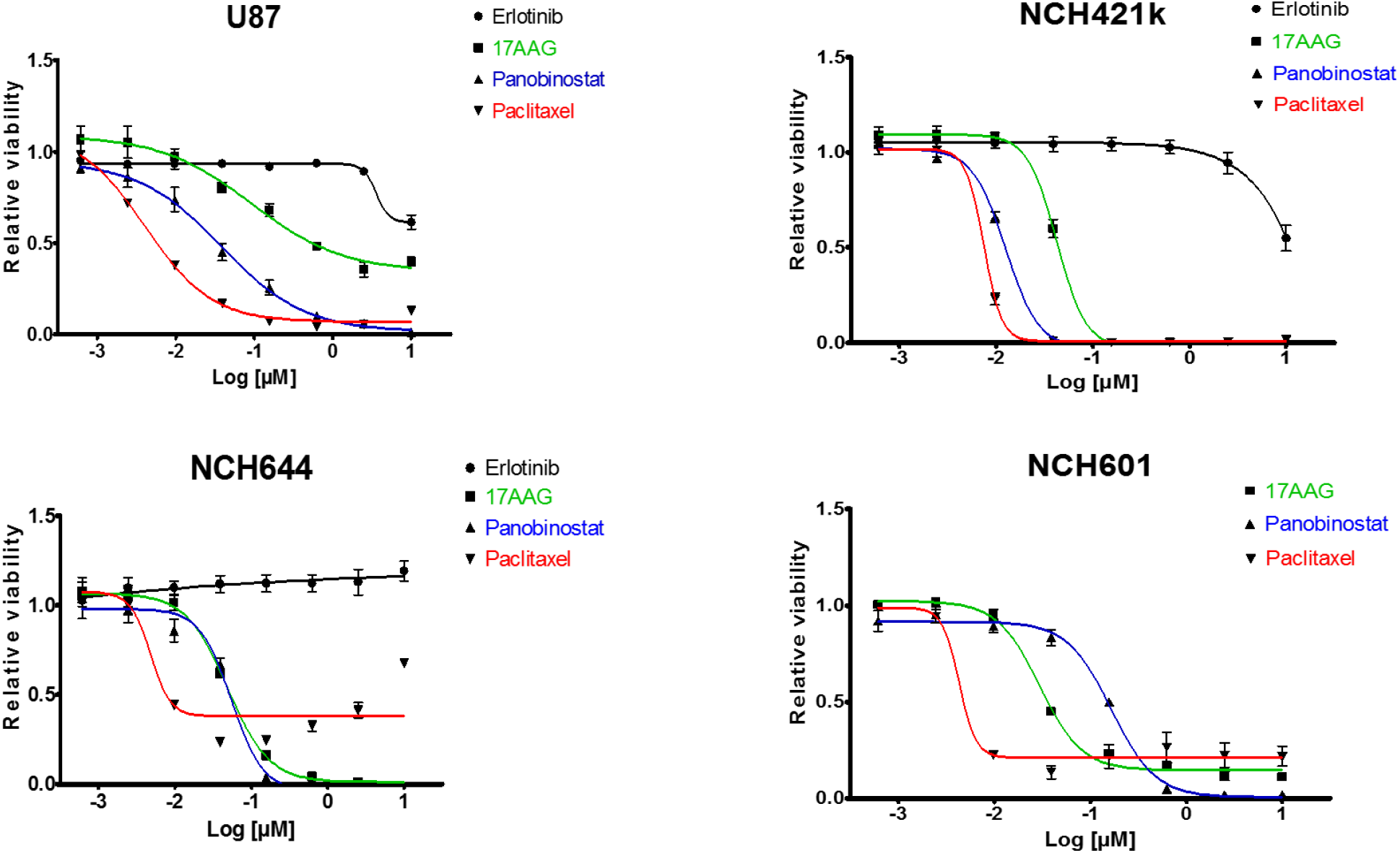
Drug response curves for the 4 drugs tested on the 4 GBM cell lines. Related to Figure 5C. Drugs were tested on each cell line in triplicates, and relative viability (vs. vehicle-treated samples) was measured for 8 drug concentration values (shown here as Log[μM]).

**Table 1.**
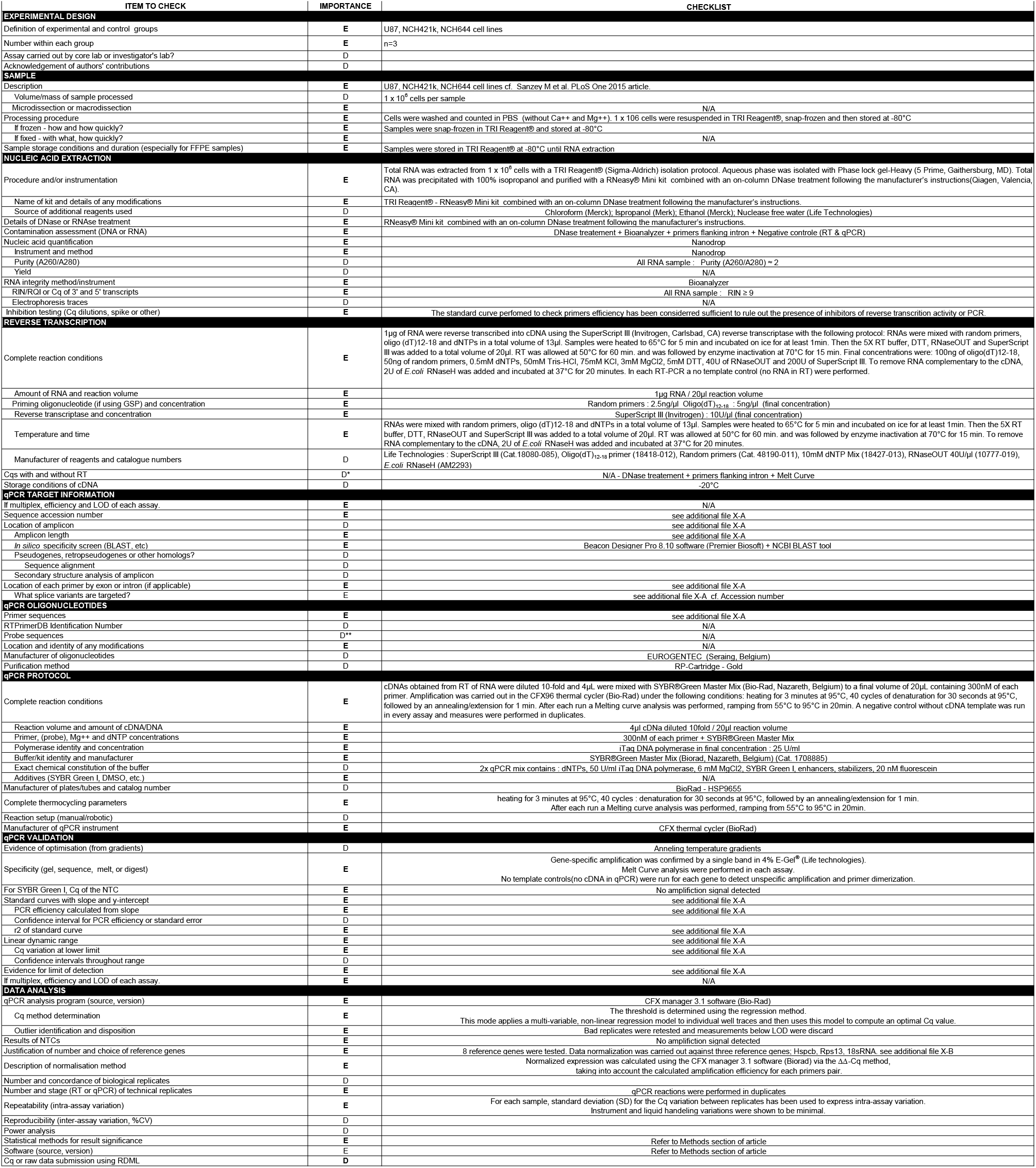
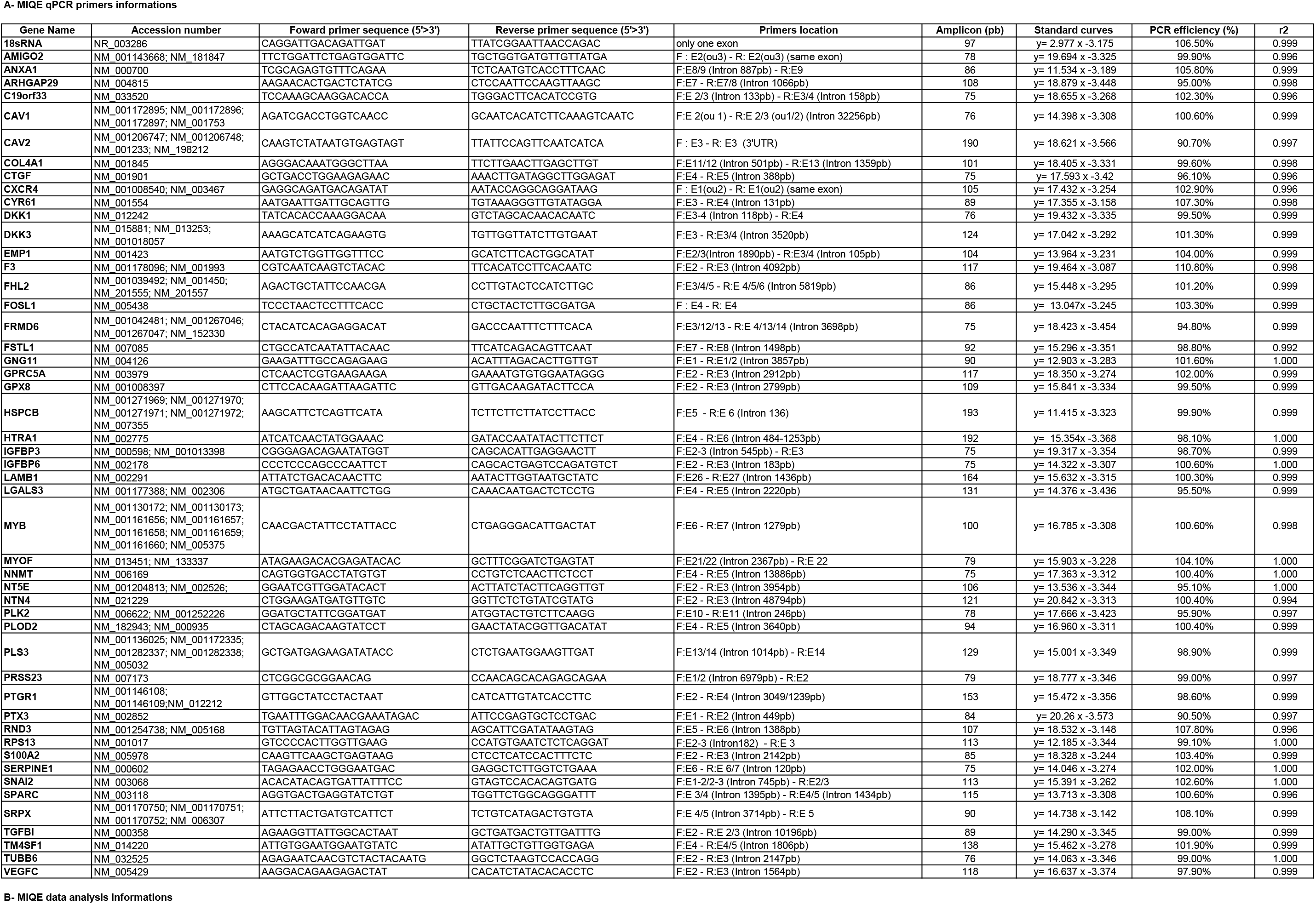
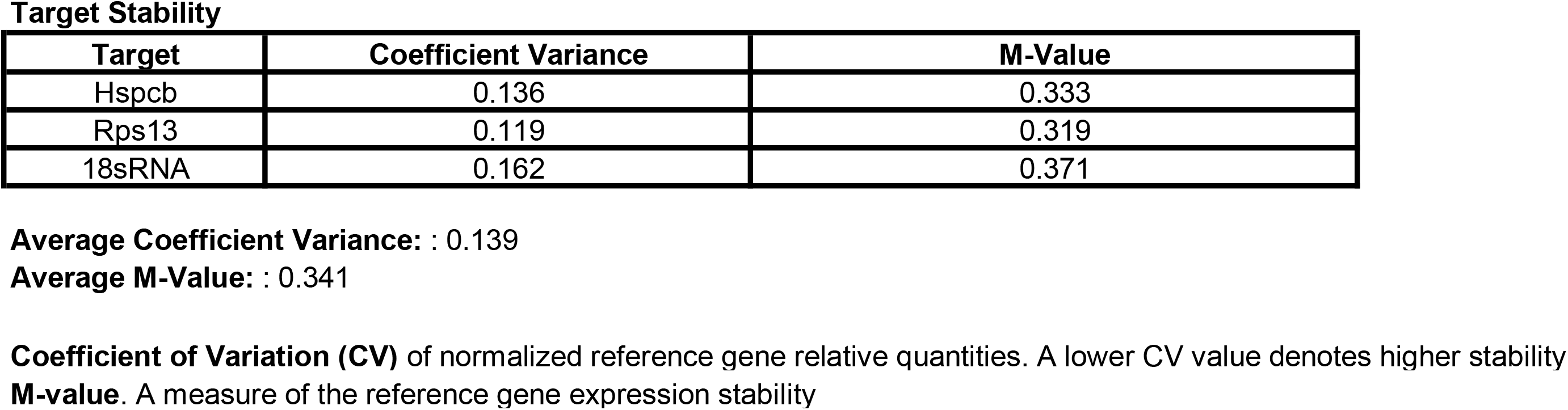
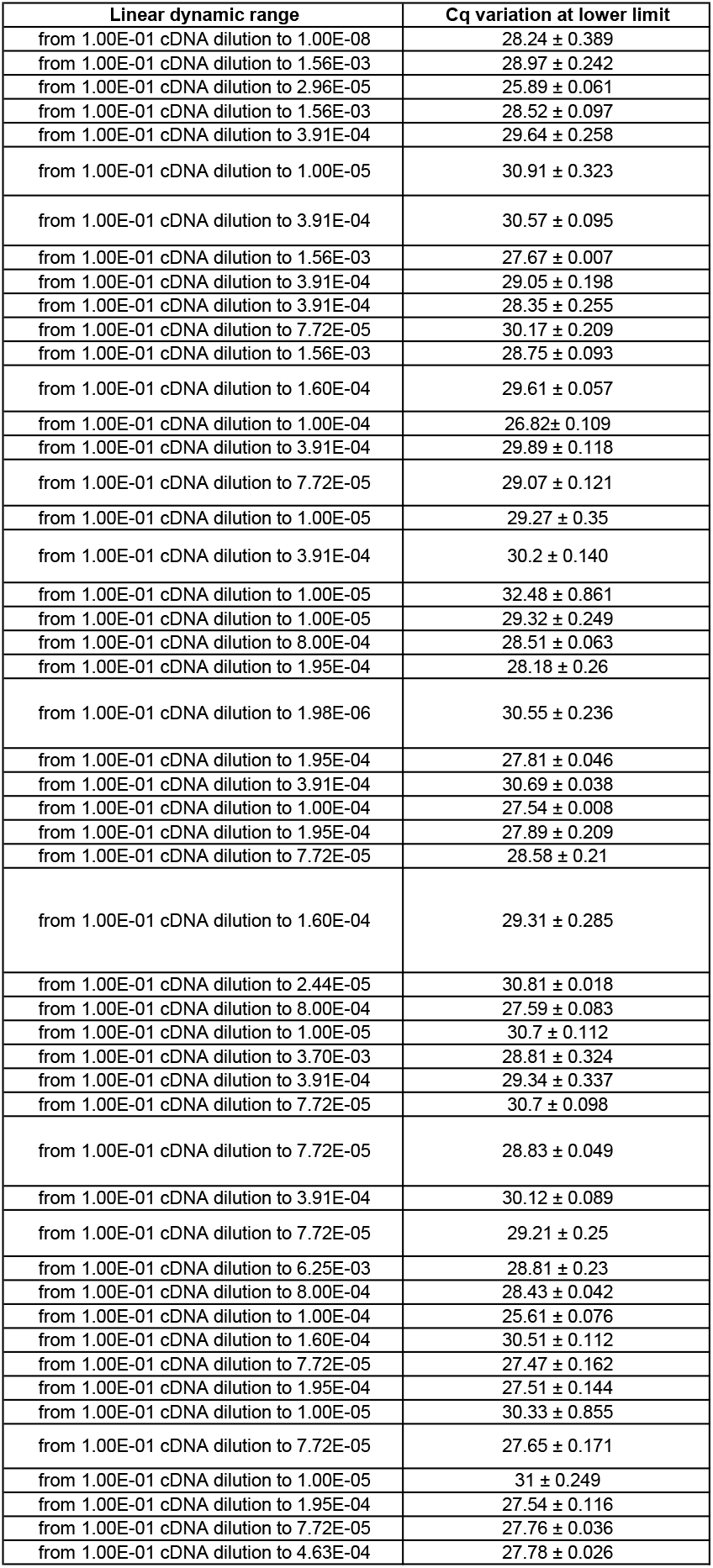
MIQE checklist for authors, reviewers and editors. All essential information (E) must be submitted with the manuscript. Desirable information (D) should be submitted if available. If using primers obtained from RTPrimerDB, information on qPCR target, oligonucleotides, protocols and validation is available from that source. *: Assessing the absence of DNA using a no RT assay is essential when first extracting RNA. Once the sample has been validated as RDNA-free, inclusion of a no-RT control is desirable, but no longer essential. **: Disclosure of the probe sequence is highly desirable and strongly encouraged. However, since not all commercial pre-designed assay vendors provide this information, it cannot be an essential requirement. Use of such assays is advised against.

